# Small volume blood-brain barrier opening in macaques with a 1 MHz ultrasound phased array

**DOI:** 10.1101/2023.03.02.530815

**Authors:** Thomas J. Manuel, Michelle K. Sigona, M. Anthony Phipps, Jiro Kusunose, Huiwen Luo, Pai-Feng Yang, Allen T. Newton, John C. Gore, William Grissom, Li Min Chen, Charles F. Caskey

**Affiliations:** Vanderbilt University, Nashville TN, USA; Vanderbilt University Medical Center, Nashville, TN, USA; Vanderbilt University Institute of Imaging Science, Nashville, TN, USA

## Abstract

Focused ultrasound blood-brain barrier (BBB) opening is a promising tool for targeted delivery of therapeutic agents into the brain. The volume of opening determines the extent of therapeutic administration and sets a lower bound on the size of targets which can be selectively treated. We tested a custom 1 MHz array transducer optimized for cortical regions in the macaque brain with the goal of achieving small volume openings. We integrated this device into a magnetic resonance image guided focused ultrasound system and demonstrated twelve instances of small volume BBB opening with average opening volumes of 59 ± 37 mm^3^ and 184 ± 2 mm^3^ in cortical and subcortical targets, respectively. We developed real-time cavitation monitoring using a passive cavitation detector embedded in the array and characterized its performance on a bench-top flow phantom mimicking transcranial BBB opening procedures. We monitored cavitation during *in-vivo* procedures and compared cavitation metrics against opening volumes and safety outcomes measured with FLAIR and susceptibility weighted MR imaging. Our findings show small BBB opening at cortical targets in macaques and characterize the safe pressure range for 1 MHz BBB opening. Additionally, we used subject-specific simulations to investigate variance in measured opening volumes and found high correlation (R^2^ = 0.8577) between simulation predictions and observed measurements. Simulations suggest the threshold for 1 MHz BBB opening was 0.53 MPa. This system enables BBB opening for drug delivery and gene therapy to be targeted to more specific brain regions.

## Introduction

A physiological barrier exists between the parenchyma and vasculature of the brain, blocking the transport of pathogens, neurotoxic plasma components, and blood cells (Sweeney et al. 2019). Vessels in the brain develop a continuous endothelial cell membrane sealed by tight junction structures and highly selective transport systems which account for this blood-brain barrier (BBB) (Armulik et al. 2010). While critical for regulating transport between the brain and the broader circulatory system, the BBB creates a major hurdle in the treatment of neurological disorders (Abbott 2005). Any candidate therapeutic agent faces the issue of low transport into brain tissues unless it is smaller than 400 Da and forms fewer than 8 hydrogen bonds (Pardridge 2012). These chemical properties exclude most small molecule drugs and all large molecule drugs with some exceptions such as receptor-mediated (Lajoie and Shusta 2015) and Trojan horse (Pardridge 2006) vehicles, or vasodilation induced by e.g. mannitol infusion (Rapoport 2000). These obstacles remain barriers in the treatment of Alzheimer’s disease, Parkinson’s disease, glioblastoma, and other neurological disorders.

Focused ultrasound (FUS) in combination with microbubbles can reversibly open the BBB noninvasively in focal brain locations and has been explored for many applications where localized drug delivery is desired. Circulating microbubbles interacting with the acoustic focus temporarily increase the permeability of the BBB through a mechanical interaction (Hynynen et al. 2001). FUS BBB opening is sufficient for transport of 70 kDa molecules (Choi et al. 2010) and magnetic resonance imaging (MRI) contrast agents of hydrodynamic diameters up to 65 nm (Marty et al. 2012). This opening size extends brain therapeutic options to include viral vectors (H. Li et al. 2021; Lin et al. 2016; Felix et al. 2021), nanoparticles (Ohta et al. 2020), neurotrophic factors (Samiotaki et al. 2015), and antibodies (Kinoshita et al. 2006). Transport of even larger molecules (2,000 kDa) has been achieved but may pose risk of permanent damage (Chen and Konofagou 2014). Safety can be improved by monitoring bubble activity during application of therapy (O’Reilly and Hynynen 2012). Safe FUS BBB opening without edema or hemorrhage has been demonstrated in macaque monkeys (Pouliopoulos et al. 2021) (N. McDannold et al. 2012) and humans (Lipsman et al. 2018) (Carpentier et al. 2016). Edema has been reported at lower pressures than those which cause red blood cell extravasation, but typically resolves within one week (Downs, Buch, Sierra, et al. 2015; Downs, Buch, Karakatsani, et al. 2015).

Effective and safe BBB opening requires accurate *in situ* pressure estimation. This is challenging in transcranial macaque procedures where pressure delivery varies across targets and skull incidence angle at magnitudes affecting BBB opening outcomes (Karakatsani et al. 2017). Most studies have used cavitation monitoring for *in* vivo pressure feedback after it was shown that harmonic and broadband emissions are linked to outcomes (Tung et al. 2010) and can be used to reduce the occurrence of hemorrhage and edema in rats (O’Reilly and Hynynen 2012). Many successful implementations of cavitation monitoring have been demonstrated in small animal models (Sun et al. 2017; Chien et al. 2021), and stable cavitation correlates with opening outcomes in these models (Sun et al. 2015). However, precise pressure delivery in small animals in not as much of a challenge compared to large animals and humans and has been effectively implemented without cavitation monitoring (Magnin et al. 2015; Choi et al. 2007). Higher skull attenuation in macaque and human skulls (which increases with sonication frequency) exaggerate the challenges associated with cavitation monitoring (Wu et al. 2014). Inspired by prior work focused on skull effects, we adopted a cavitation monitoring strategy which captures baseline spectra for all candidate amplitudes prior to therapy (H. A. Kamimura et al. 2019). This provides a means to adaptively change pressure during therapies based on spectral content while removing the effects of reflected sound with changing transmit amplitude. During all therapies, we plotted stable and inertial cavitation metrics as done in similar systems along with an updating spectrogram and line plot of the latest spectrum (Marquet et al. 2014; Pouliopoulos et al. 2021; Chien et al. 2021; Novell et al. 2020).

In glioblastoma treatment, large BBB opening volumes on the order of 1-2 cm^3^ are desirable to match tumor size (Idbaih et al. 2019). However other applications require small opening volumes to match the anatomical target’s size. The transducer tested in this work is designed for gene therapy at the frontal-eye field (FEF) and benefits from restricting the gene delivery to the FEF alone. The macaque FEF is approximately 10 mm in depth at its largest cross-section, 3 to 9 mm wide and has a total volume of 211 mm^3^ (Jung et al. 2021). Most BBB openings demonstrated in macaques have used single element transducers ranging from 200 to 500 kHz which are well suited for transmission through the skull but have spot sizes much larger than the FEF. The transducer in this study has a higher central frequency of 1 MHz and a spot size of 1.9 × 1.9 × 9.5 mm in free field when steered inward 10 mm, making it well-suited for this target. We described the design and characterization of this transducer in a prior publication (Manuel, Phipps, and Caskey 2022).

The goal of this work is to investigate the capabilities of a transducer optimized for small volume BBB opening in macaques, including developing cavitation-based feedback to identify the pressure range which opens the BBB without causing edema or hemorrhage. We characterize the performance of a cavitation monitoring system in *in vivo* and benchtop scenarios, and then applied the system to open the macaque BBB and quantify the opening volumes achieved at different cortical and subcortical targets, including the FEF. Acoustic simulations were performed to better understand the variance in opening volumes occurring during these sonications. Our study is the first to characterize ultrasound-based BBB opening with 1 MHz through intact macaque skulls at cortical and subcortical targets, outlining a safe pressure range and highlighting challenges with cavitation monitoring through the skull at high frequencies.

## Methods

### Transducer specifications

A custom built transducer was used for therapies (Imasonic, Besancon, France). The transducer is a 1 MHz array with a total diameter of 58 mm, a focal length of 53.2 mm (f-number as 0.92), and 128 3.5 mm diameter transmit elements distributed along a Fermat spiral. The transducer features one central receive element for cavitation monitoring with peak sensitivity at 2 MHz and an active diameter of 3.5 mm. The transducer’s un-steered spot size is 2.2 × 2.0 × 13.5 mm (31 mm^3^) and decreases with steering towards the transducer. We leveraged this to reduce opening volume, using an inward steering of 10 mm at cortical targets. At 10 mm, the free-field spot size reduces to 1.9 × 1.9 × 9.5 mm (17.95 mm^3^). We expect the focus to broaden due to aberration in *in vivo* procedures. Simulations predict a transcranial spot size of 2.5 ± 0.3 mm lateral and 9.5 ± 1.0 mm axial (30 ± 8 mm^3^) with conditions used during our cortical therapies (10 mm inward steering and no aberration correction). Transcranial transmission to the frontal eye field was 27 ± 6% in design simulations.

We 3D-printed a custom transducer cone (Stratasys, Rehovot, Israel) which paired with a neoprene membrane bound by rubber o-rings and a degassing circuit to provide coupling to the head during procedures. This setup is described in more detail here (Manuel, Phipps, and Caskey 2022). The transducer was powered by a 10-watt-per-channel, 128-channel generator (Image Guided Therapy, Pessac, France) which was impedance-matched to the transducer via a matching network.

### Animal protocol

The ultrasound procedures were performed in two adult macaques (one male, one female) with a minimum of two weeks between sessions in the same macaque. During the procedures animals were initially anesthetized with ketamine hydrochloride (10 mg/kg) and then anesthetized with isoflurane (1.0-1.5%) delivered over medical air. Medical air was used over oxygen delivery because oxygen has been shown to reduce the half-life of circulating microbubbles compared to medical air (Mullin et al. 2011). 2.5% dextrose in saline solution was infused intravenously (3 ml/kg/h) to prevent dehydration. Artificial ventilation was used during the procedure. Animals were placed in a custom stereotactic frame with ear bars, eye, and mouthpieces to secure the head (figure 1). A circulating water blanket informed by a rectal temperature probe was used to maintain body heat between 37.5°C and 38.5°C. Respiration pattern, heart rate, end-tidal CO_2_ (24-32 mmHg; SurgiVet), and peripheral capillary oxygen saturation (SpO_2_; Nonin) were monitored and maintained during the procedure. All procedures were conducted in accordance with National Institutes of Health guidelines and were approved by the Institutional Animal Care and Use Committee of Vanderbilt University. All MR images were acquired using two MRI surface coils placed on opposite sides of the head (MRI methods are described below).

**Figure 1.**
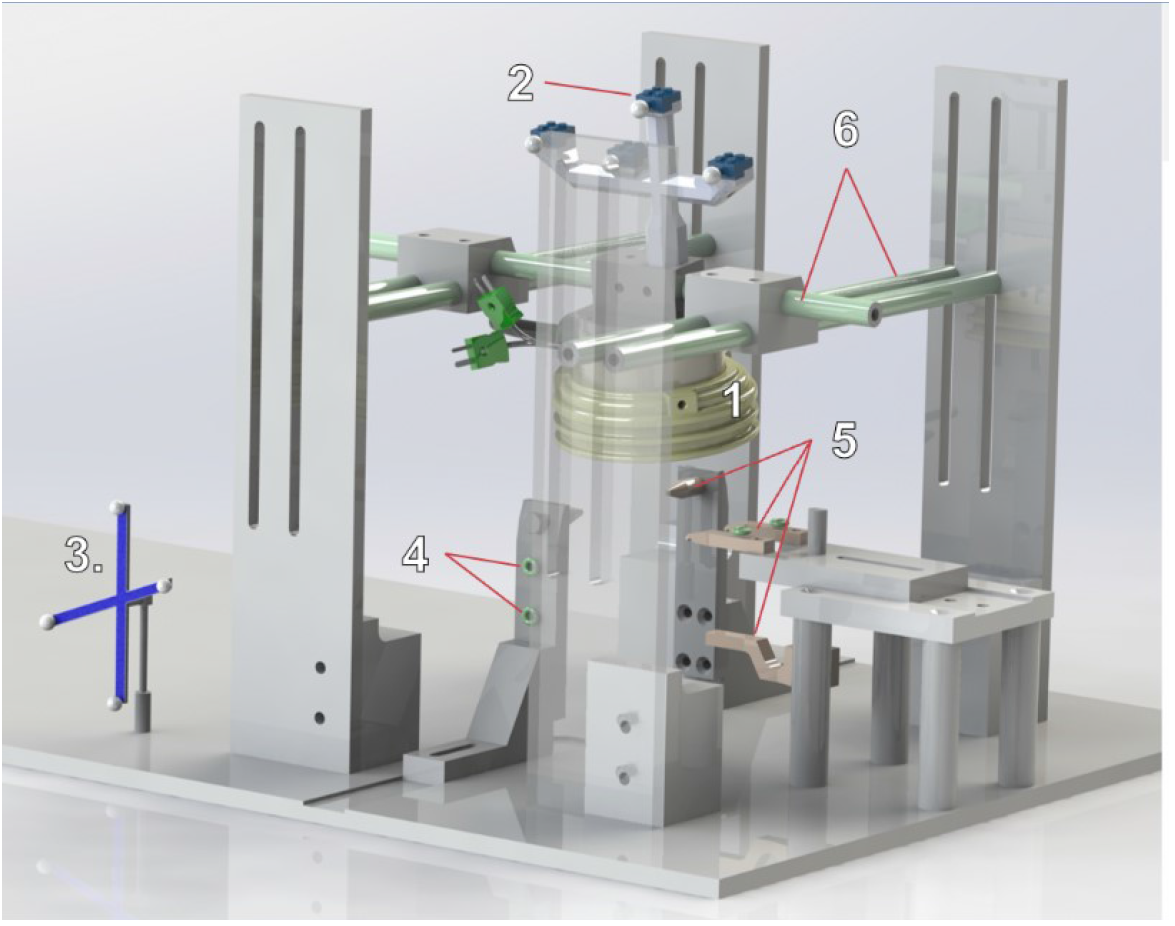
Mechanical and optical tracking setup used during in vivo BBB opening procedures (water bag for coupling not shown).

### Neuronavigation

The transducer was mechanically moved with the custom stereotactic frame guided by optical tracking (Figure 1) as described previously (Phipps et al. 2019; Chaplin et al. 2019). A Polaris Vicra optical tracking system (Northern Digital Inc., Ontario, Canda) was used for tracking. An MRI compatible rigid body tracker was fixed to an NHP table which also held the stereotactic frame. The fixed tracker was used as an optical tracking global reference to track a separate tracker fixed to the transducer and a stylus used to locate six multimodality fiducials (IZI Medical Products, Maryland, USA) distributed on the ear bars and eyepieces holding the monkey head. The fiducials were visible on T_1_-weighted images, allowing registration between MR image-space and optical tracking space which was performed using 3DSlicer’s Image Guided Therapy module (Ungi, Lasso, and Fichtinger 2016). Optical tracking was used for initial guidance of the transducer to the target by projecting a model of the focus onto the MR images. The transform used to orient the focus relative to transducer tracker was determined in a water bath experiment by measuring the location of the focus relative to the transducer’s rigid body tracker using an optically tracked hydrophone (Manuel, Phipps, and Caskey 2022). Tracking was visualized along with MR images in 3DSlicer (http://www.slicer.org/) (Fedorov et al. 2012). Magnetic resonance acoustic radiation force imaging (MR-ARFI) was used to confirm the position of the focus relative to brain anatomy prior to each therapy (Nathan McDannold and Maier 2008) and inform electronic steering of the focus if necessary. The MR-ARFI pulse sequence has been previously described (Phipps et al. 2019). Therapies were performed in a 3 Tesla MRI scanner (Phillips Healthcare, Elition X, Amsterdam, Netherlands).

During procedures, we first selected a target on a T_1_-weighted image. Next, we oriented the transducer such that the optically tracked focus was close to the target of interest. We then moved the monkey, frame, and transducer into the MRI bore and collected MR-ARFI to measure the displacement generated by the acoustic focus. If the displacement overlapped with grey matter near our target of interest, we moved forward with therapy. If not, we steered the beam electronically while collecting MR-ARFI images until we measured displacement in grey matter at our target. In several cases, particularly at cortical targets, the displacement images were insufficient for focus identification. In these cases, we acquired an additional MR-ARFI with the beam electronically steered 1 cm away from the skull. In those cases, observing displacement in regions beneath a grey matter zone at our target confirmed our positioning.

### Acoustic therapy

All therapies used 10 ms, 1 MHz pulses repeated at 2 Hz for 2 minutes. Injections were intentionally slowed to take between 15 and 30s to prevent bubble collapse while traveling through the syringe needle. To achieve consistent therapy length, we terminated therapies 2 minutes following the completion of a saline flush performed to clear remaining bubbles from the injection port and catheter line. Therapies were performed at the foot of the MRI bed (∼7 feet outside the bore) to minimize the damping effect of a strong magnetic field on bubble oscillations (Yang et al. 2021). We tested peak negative pressures (PNP) between 0.4 to 1.4 MPa. Once hemorrhage was detected at 1.4 MPa, no pressures at or above that range were tested again.

Definity microbubbles (Lantheus Medical Imaging, North Billerica, MA, USA) were administered at 20 uL/kg diluted into 3 mL of saline. Following therapy, a T_1_-weighted image was collected. Gadavist (Leverkusen, North Rhine-Westphalia, Germany) was injected at a dose of 0.1 mL/kg and circulated for 5 min prior to collection of a second T_1_ -weighted image. Susceptibility-weighted images were collected to check for extravasation of red blood cells, and fluid-attenuated inversion recovery (FLAIR) images were collected to check for edema (Ho, Rojas, and Eisenberg 2012).

Assigning a single transmission value is an oversimplification because transmission changes with each subject, target, and transducer orientation. However, ascribing derated pressure estimates aids in visualization and interpretation of results. *In situ* pressure for display was estimated as 27% of free field PNP. This value was taken from design simulations of this transducer in macaques which predicted a transmission of 27 ± 6% through four monkey skulls at the FEF (Manuel, Phipps, and Caskey 2022). The transmission estimates were also informed by water tank measurements through an *ex vivo* macaque skull which gave transmissions from 15% to 40% depending on skull orientation. When positioning the transducer, we attempted to minimize the angle between the transducer and the skull to avoid the effects of angle on BBBO outcome (Karakatsani et al. 2017).

### MR-Imaging Parameters

T_1_ -weighted: 3D magnetization-prepared gradient echo sequence, TR/TE: 9.9/4.6 ms; flip angle: 8°; in plane resolution: 1 mm^2^; slice thickness: 1 mm with 0.5 mm slice overlap reconstructed to 0.5 mm isotropic voxels. Susceptibility-weighted (SWI): 3D spoiled gradient recalled echo, TR 31 ms; 4 TEs (7.2, 13, 20, 26 ms); flip angle: 17°; in plane resolution: 0.5 mm^2^; slice thickness: 1 mm with 0.5 mm slice overlap reconstructed to 0.33 × 0.33 × 0.5 mm. FLAIR: 3D inversion recovery segmented k-space; TR/TE: 4.8/0.34 ms, inversion time: 1.65 ms; flip angle: 90°; in plane resolution: 1 mm^2^; slice thickness: 1 mm with 0.5 mm slice overlap reconstructed to 0.5 mm isotropic; 2 acquisitions.

### Image processing

All pre and post gadolinium T_1_-weighted images were processed by FSL’s BET tool (Smith 2002) for brain extraction and then by FSL’s FAST segmentation algorithm (Zhang, Brady, and Smith 2001). The FAST algorithm was used for bias field correction and segmentation into tissue types. A rough crop of the head was required prior to BET for successful brain extraction. Five iterations were used for bias field correction (option -n). Following bias field correction, several of the images showed histogram differences beyond what would be expected from the presence of gadolinium alone. If unadjusted, the subtraction images generated from the pre- and post-gadolinium injection T_1_ -weighted images displayed whole-brain differences which made quantifying opening difficult. We devised a two-step process to adjust for this. First, we balanced the images by iteratively applying a scalar offset to the post-gadolinium image while minimizing the difference of tissue type thresholds output by the Otsu threshold technique (Otsu 1979). Next, we used MATLAB’s histogram matching algorithm (imhistmatch) to match the histogram of the post-gadolinium image to that of the pre-gadolinium image.

Figure 2 illustrates the processing steps applied to the percent change images. Percent change images were created using the balanced and histogram matched pre- and post-gadolinium T_1_ -weighted images. The percent change images were cropped using a cylinder mask centered on the acoustic focus of size equal to three times the free-field pressure size (full-width at half maximum) which corresponds to a cylinder of height 30 mm, diameter 9 mm, and volume 1908 mm^3^. This mask was created using 3DSlicer’s “create model” module and placed at the location with angled orientation informed by optical tracking and MR-ARFI. Cropping the percent change image reduced off-target mislabeling of BBB opening at noisy regions and blood vessels in the image far enough away from the focus to rule out as BBB opening. For white matter/grey matter quantification, the cropped percent change image was registered to a tissue type atlas for the corresponding therapy. Grey/White/CSF atlases were created from FSL’s FAST segmentation algorithm. The caudate and putamen segmentations required manual correction in several therapies where subcortical grey matter was mislabeled as white matter. Opening volumes are reported using enhancement thresholds of 10%, 20% and 30%.

**Figure 2.**
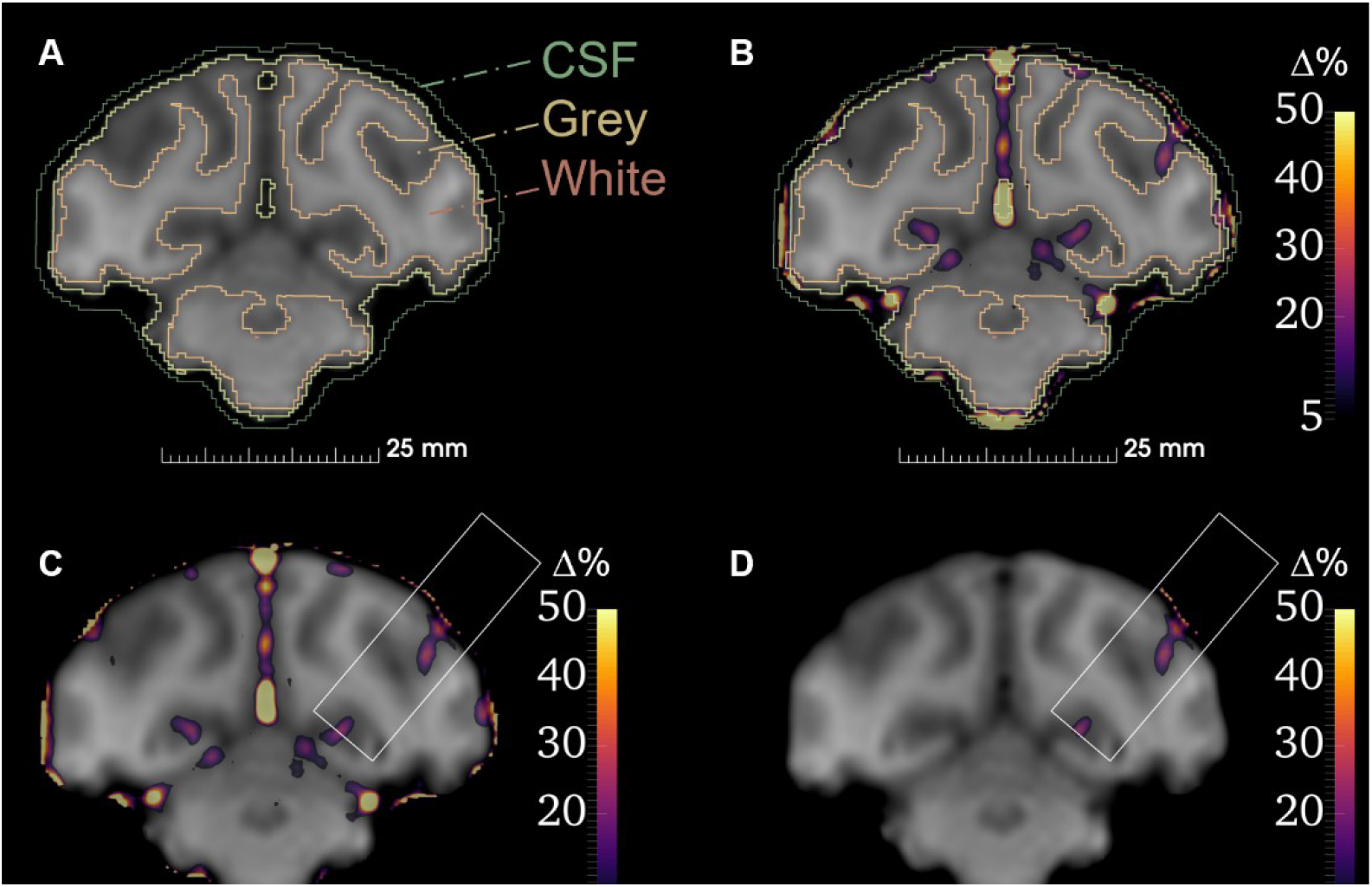
A portion of the image processing pipeline used to quantify opening volumes. A) segmentation into tissue types; B) overlay of full percentage change image onto segmentation; C) Cylinder mask used to crop around targeted region; D) Final percent change image used to quantify opening.

### Acoustic simulations of BBB opening therapies

We simulated the conditions of each therapy with the aim of comparing predicted opening volumes to measured opening volumes. Simulations ran in k-Wave used a 0.2 mm isotropic grid (7.5 points per wave, 1 MHz, water) (Treeby and Cox 2010). Computed tomography scans captured at 0.6 × 0.6 × 0.8 mm and upsampled to 0.2 mm isotropic were used to estimate speed of sound and density from Hounsfield units (HU). Density of voxels were estimated from HU by referencing the HU of water and air (Connor, Clement, and Hynynen 2002; Pichardo, Sin, and Hynynen 2011) given by:

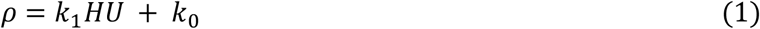

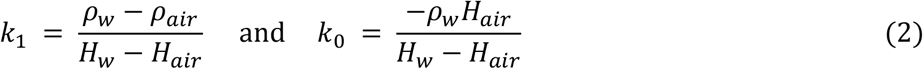

with ρas density, ρ_*w*_density of water, ρ_*air*_density of air, *H*_*w*_HU of water, and *H*_*air*_HU of air. Speed of sound was estimated by first estimating bone porosity *φ*:

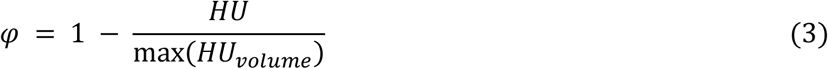

and then estimating the speed of sound (Aubry et al. 2003):

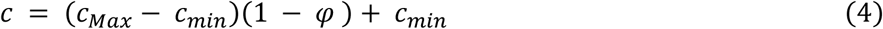

Values used were *c*_*Max*_3100 m/s, *c*_*Min*_1480 m/s, *ρ*_*w*_997 kg/m^3^, *ρ*_*air*_1.225 kg/m^3^, *H*_*w*_0, *H*_*air*_-1000. To model absorption, a value of 0.2 dB/MHz/cm was used for water voxels, 8.1 dB/MHz/cm for skull (Pinton et al. 2012) and 0.8 dB/MHz/cm for soft tissue (brain and muscle). Soft tissue was assigned a speed of sound 1560 m/s.

All registration was performed in 3DSlicer. Transducers were placed in the therapy orientation by creating a model which could be positioned by the optical tracking transforms already used in therapies for image guidance. The creation of these models and workflow between 3DSlicer and k-Wave was previously described in (Manuel, Phipps, and Caskey 2022). An additional translational transform was applied if an offset was observed between the predicted focus location from optical tracking and that measured by MR-ARFI. For therapies that used electronic steering, the phases used for steering were input into simulations to produce the equivalent beam steering. In *in vivo* therapies, pressure was varied over time based on cavitation signal. Simulation pressures were matched to the average of the free-field pressures of the corresponding therapy. This required calibration simulations for each therapy to find the corresponding amplitude value to use for the k-Wave source which would produce the desired free-field pressure for the given steering coordinate. For each therapy, three free-field simulations were run at increasing amplitude values. From these a linear fit was used to determine the slope between focus pressure and source amplitude for the given steering coordinate. From this slope, the appropriate amplitude value could be calculated to produce a free-field pressure matching that of the average free-field pressure used in the therapy. Simulation outputs were co-registered with the tissue type atlases generated for *in vivo* data processing. Predicted opening volume was taken as the volume of grey matter voxels subjected to a pressure threshold. Simulation outputs were analyzed with pressure thresholds varying from 200 kPa to 1 MPa at 25 kPa increments.

### Cavitation water tank measurements

Figure 3 shows a flow phantom and setup designed to mimic *in vivo* therapy conditions to enable benchtop testing of the cavitation monitoring system. The phantom used 4 mm ID soft PVC plastic tubing at the point of the acoustic focus. We used a flow circuit powered by a 12V variable speed pump driven by a variable power supply to circulate the microbubble solution. We tuned the voltage on the power supply to achieve a 1 cm/s flow velocity. Velocity was measured by introducing a visible air bubble into circulation and timing its traversal through a known length of tubing. We matched estimated *in vivo* microbubble concentrations by converting the microbubble dose (20 µl/kg) and the average blood volume in adult macaques (60 ml/kg) which gives a ratio of 20 µl microbubble solution per 60 ml of water to use in the flow phantom (Bender 1955). For water tank measurements we used in-house manufactured microbubbles following a previously described method (Singh et al. 2022). Stirring was required to keep the bubble solution mixed during experiments and was achieved by placing a reservoir on a magnetic stirrer. The bubble tube was held by a 3D-printed y-shaped adapter mounted to a 3-axis stage. The tube was aligned with the acoustic focus while filled with air by maximizing the amplitude of the reflection off the tube recorded at the receive element. Water tank measurements through skull were achieved using an *ex vivo* monkey skull cap which was degassed for 24 hours at -95 kPa while submerged in a cylindrical vacuum chamber (Abbess Instruments, Holliston, MA, USA). The transmission through this skull fragment was between 15 and 40% depending on skull orientation (measured using a ceramic needle hydrophone). Therefore, to attain an estimate of transmission for the specific orientation during the flow experiment, we captured a 5-cycle pulse-echo off the flow tube filled with air with and without the skull present by transmitting with the therapy elements and receiving with the PCD. With the skull present, the receive amplitude decreased by 87.5%. The one-way transmission loss was estimated from this two-way measurement as 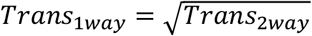 or 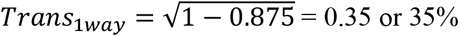.

**Figure 3.**
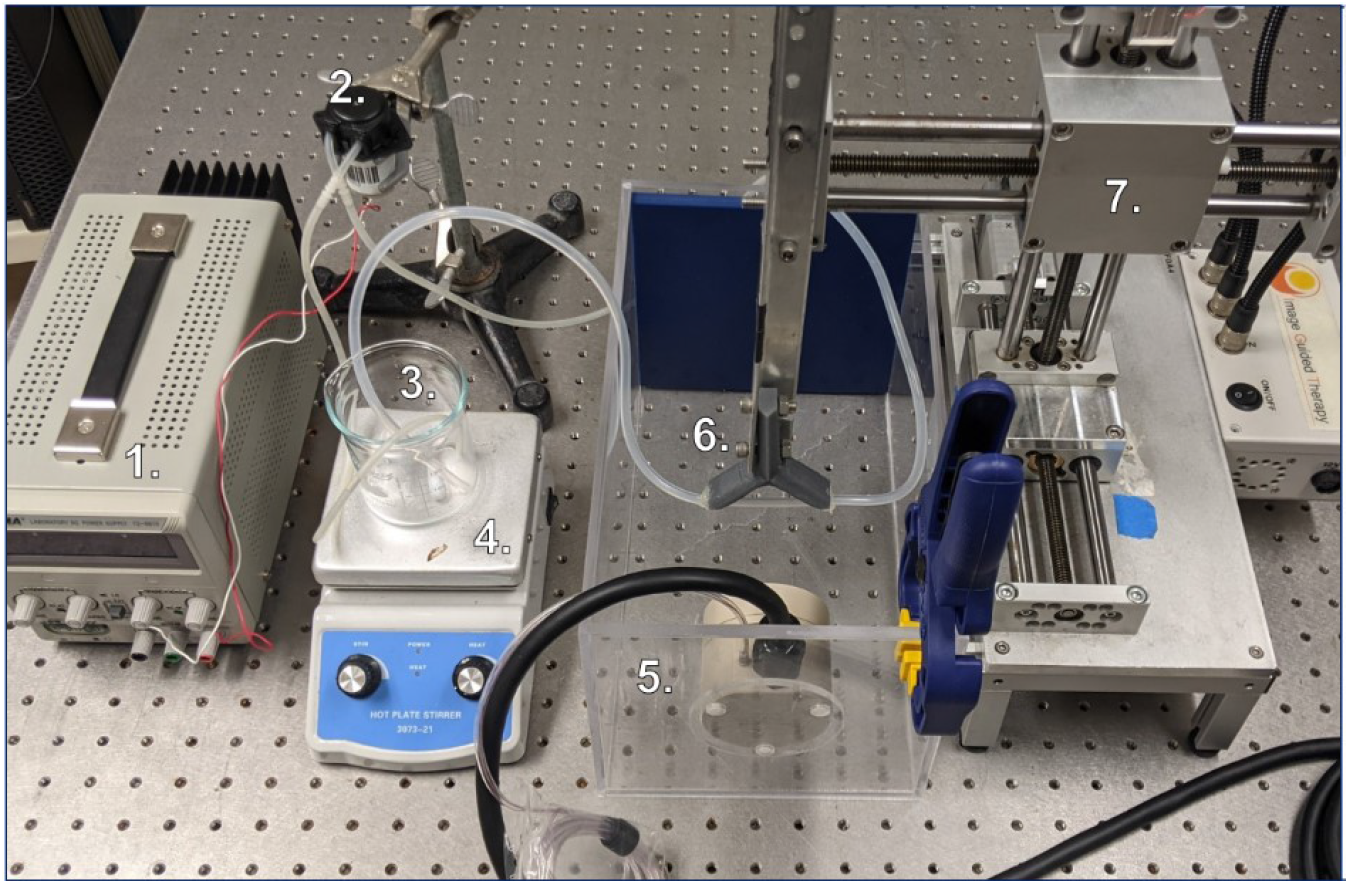
The setup used for water tank microbubble flow phantom measurements to develop and validate the cavitation monitoring system.

### Cavitation monitoring software

We built a Python application to monitor cavitation. The design of the monitoring software emphasized the ability to visualize cavitation signals in real time, change therapy amplitudes, and present metrics which inform if *in situ* pressure reaches unsafe levels. This application communicates between a Picoscope ps5000a (Pico Technology, Cambridgeshire, U.K.) mounted within our generator cabinet, the SDK provided by Image Guided Therapy for generator control, and user inputs given through a graphical user interface (Tkinter). The sample rate was fixed at 62.5 mega samples per second and 500 µs captures. One capture was acquired after a 50 µs delay following each therapy pulse trigger (2 Hz). Prior to therapy, the software allows the user to define a range of candidate amplitudes and an electronic steering coordinate. Prior to bubble injection, the software acquires baseline captures for each candidate amplitude. Twenty captures per baseline are acquired and averaged in the frequency domain. These baselines enable dynamic baseline subtracting i.e., subtracting the baseline which corresponds to the current therapy amplitude. This aids in decoupling effects from reflection amplitude changes and microbubble emission changes and was first adopted by Kamimura and colleagues (H. A. S. Kamimura et al. 2016).

During therapy the software displays four windows: a spectrogram; a 2D line plot of the latest spectrogram column; plots of stable cavitation and inertial cavitation metrics versus time; and a control panel to start and stop therapy, change amplitude, and save data. For each therapy pulse, a column *n* in the spectrogram was calculated as

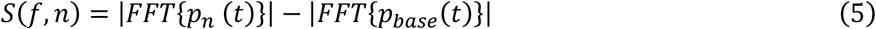

where *S* is the spectrogram with rows in frequency space *f* and columns *n*corresponding to the pulse/acquisition number. *p*_*n*_*(t)* is the latest acquired pressure-time series and *p*_*base*_*(t)*is a baseline pressure-time series from a pulse with the same current transmit amplitude. *FFT* is the discrete fast Fourier transform.

Our PCD has a central frequency of 2 MHz and detects harmonics from 0.5 MHz to 3 MHz in water tank measurements with a flow phantom and no skull present. Using this as an assessment of effective bandwidth, we limited frequency analysis within these bounds. Stable cavitation and inertial cavitation metrics are calculated by masking relevant frequency bands in *S*. The stable cavitation mask, *SCM*, corresponded to ±10 kHz bands surrounding 0.5, 1.5, 2, 2.5 and 3 MHz. 10 kHz bands were chosen based on prior work (Pouliopoulos et al. 2021). The inertial cavitation mask, *ICM*, spanned 1.1 MHz to 2.9 MHz excluding portions overlapping with the stable cavitation mask. Inertial cavitation dose, *ICD*(*n*), and stable cavitation dose, *SCD*(*n*), metrics were calculated for the n^th^ pulse by:

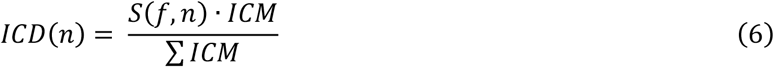

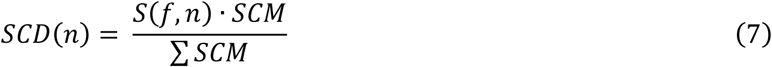

Here. represents the dot product. We divide by the sum of the masks to normalize values such that *ICD*(*n*) and *SCD*(*n*) are not scaled by the different number of frequency points in both. Dynamic colormap windowing was necessary to visualize the spectrogram with sufficient contrast. We calculated a new lower and upper bound for the colormap with each pulse. The lower bound was equal to the average of the latest spectrogram column. The upper bound was equal to the average plus 3 standard deviations of the spectrogram column. Rather than automate pressure changes, we manually adjusted pressure during therapies because the signal amplitudes and qualities varied greatly from target to target.

### Overview of Therapies

We performed twelve therapies in two macaques (one male, one female) over the course of five separate days. Targets were selected to investigate both BBB opening capabilities throughout the brain as well as BBB opening capability at the frontal eye field (FEF), the target this transducer was optimized for in the design stage (Manuel, Phipps, and Caskey 2022). Nine of the twelve therapy targets were cortical, with four performed at the FEF. The other three targets were subcortical (two putamen, one caudate). A range of pressures were attempted with mean *in situ* pressures ranging from 0.4 to 1.4 MPa.

## Results

### Opening volume

Figure 4 shows percent change images generated from pre and post therapy T_1_-weighted images overlaid on T_1_-weighted anatomical images for three therapies from each target subgroup (subcortical, cortical, FEF only). Opening regions largely follow gray matter topology and vary in shape and size. Subcortical therapies resulted in enhancement at several cortical regions along the path to the transducer focus despite being outside the spot size of the transducer. Subcortical targets used the geometric focus of the transducer, rather than steered inward 10 mm. The volume of the acoustic geometric focus is 1.7 times larger than when steered inward and resulted in larger opening volumes than achieved at cortical targets. For some cortical targets, contrast enhancement can be seen in the subarachnoid space above the outermost grey matter as well as in grey matter regions, matching prior clinical results (Carpentier et al. 2016).

**Figure 4.**
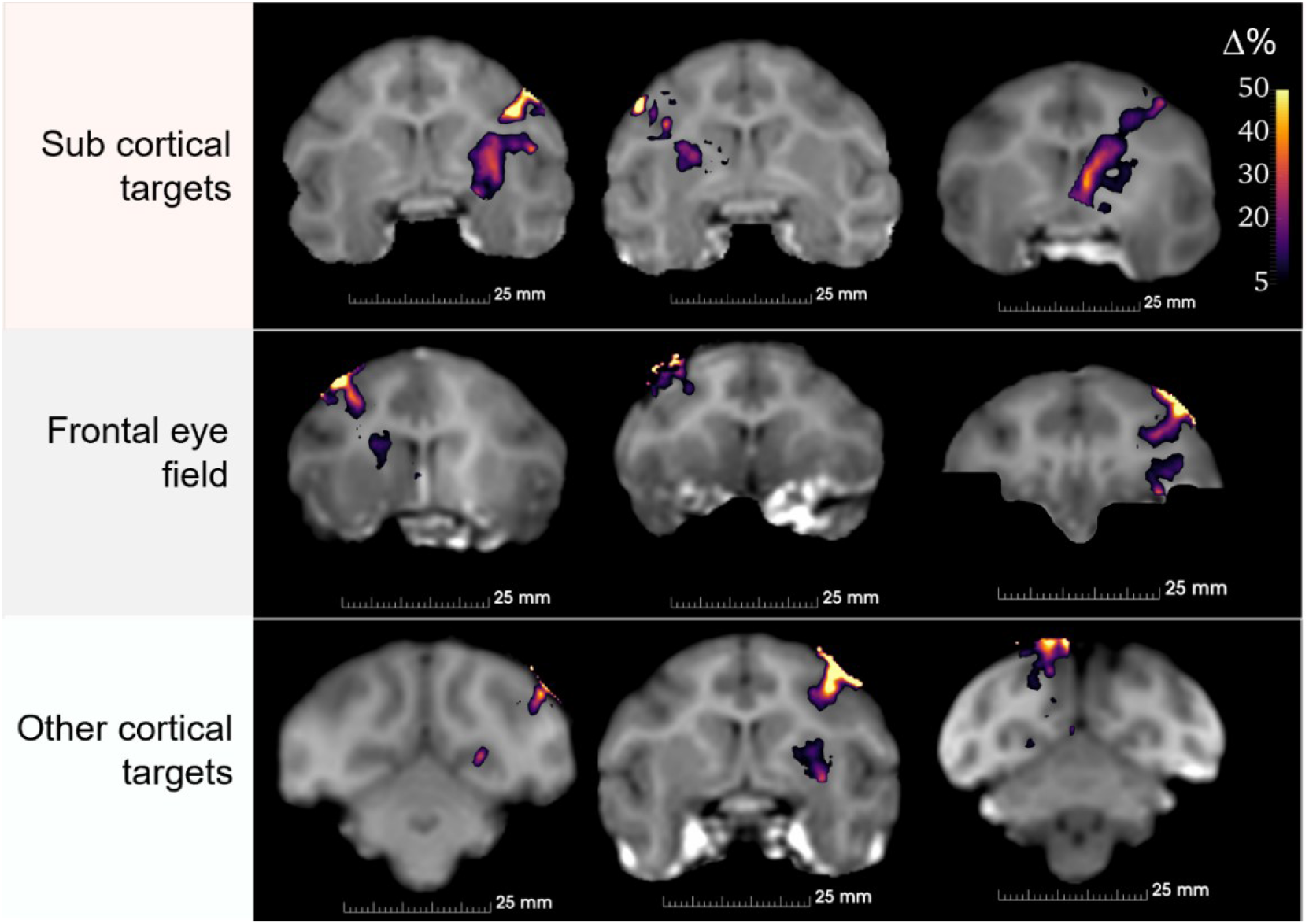
Percentage change images for nine BBB opening therapies separated into columns based on target groups. All colormaps match that shown in the top right. Subtraction images are overlaid on T1-weighted images from the same therapy.

The average tissue volume (grey + white matter) which experienced greater than 10% enhancement in post gadolinium T_1_-weighted images was 103 ± 101 mm^3^. The opening volume at subcortical targets was 184 ± 82 mm^3^. The opening volume of cortical targets was 59 ± 7.3 mm^3^. Figure 5 panel A shows a breakdown of contrast enhancement between grey and white matter tissues separated into cortical and subcortical targets. 88% of opening was in grey matter when considering all targets. Figure 5 also displays volumetric enhancement at thresholds of 20% and 30%. The enhanced volume was 31 ± 9 mm^3^ and 9 ± 10 mm^3^ for 20% and 30% thresholds respectively. 97% of enhancement greater than 20% was in grey matter (excluding subarachnoid space, not included in analysis). We compared opening volumes against mean and maximum therapy pressure (figure 5, panel B). Opening volume increased with increased mean pressure (R^2^ = 0.55) and increased maximum pressure (R^2^ = 0.53).

**Figure 5.**
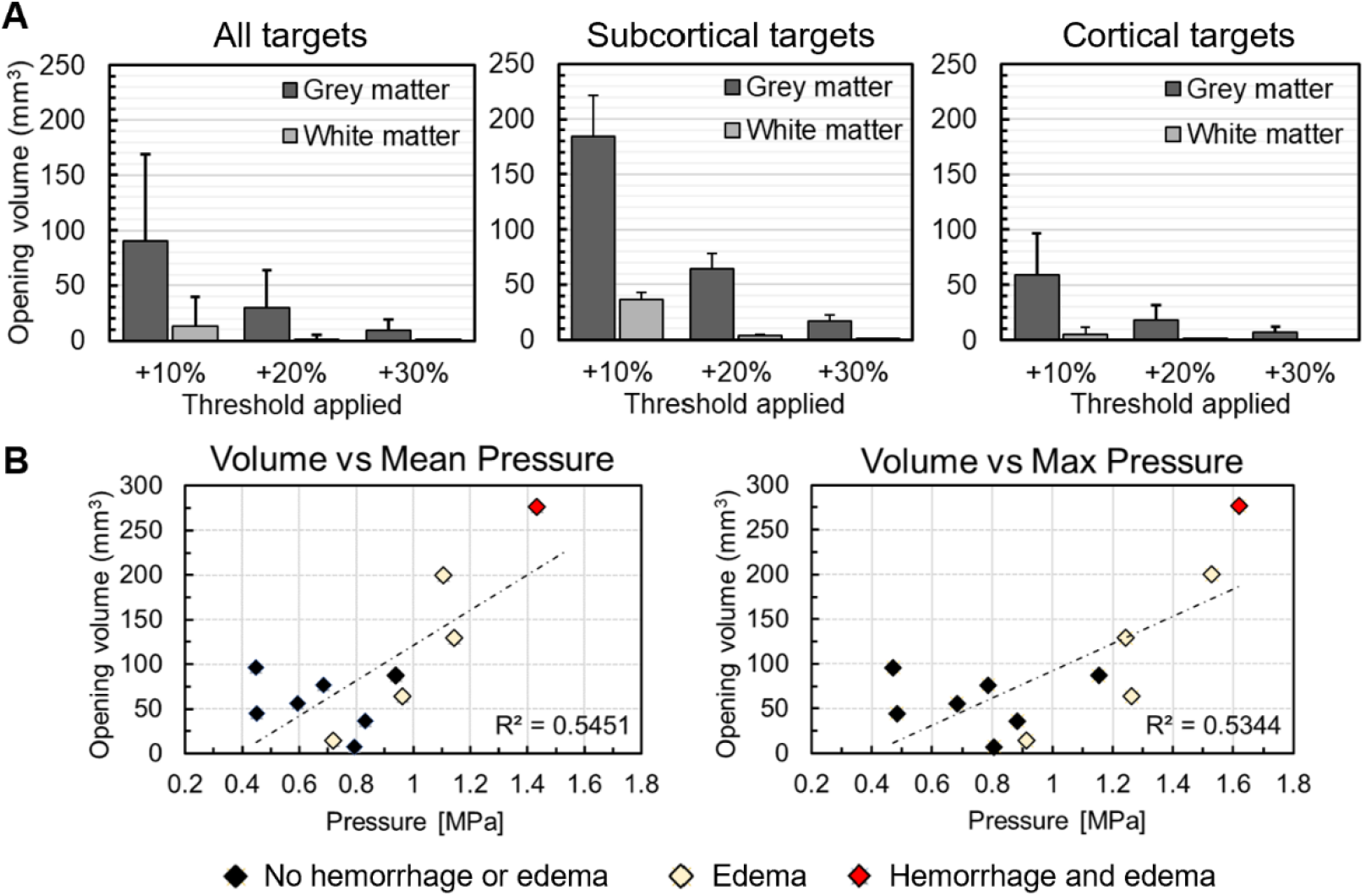
Opening volume results. A) Opening volume for all targets, subcortical targets, and cortical targets in grey and white matter. For each group opening volume is presented using a 10%, 20%, and 30% enhancement threshold. B) Opening volume compared to mean and maximum pressures used during therapies. Data points are color coded to indicate if edema or hemorrhage were detected.

### Simulating in vivo therapy conditions

Simulated opening volumes were correlated with measured opening volumes (R^2^ = 0.8577) (figure 6, panels A, B). Average simulated transmission was 29.8 ± 6.3 % which is close to our a priori estimate of 27 ± 6 % transmission. By computing resulting simulated opening volumes in grey matter using pressure thresholds from 200 kPa to 1 MPa, the pressure threshold which most closely matched measurements was 0.53 MPa (figure 6, panel C). Using 0.53 MPa as the threshold resulted in an average difference between simulation and measurement across therapies of 35 ± 23 mm^3^. The simulated opening volumes were 25.4 ± 20.3 mm^3^ and 171.9 ± 7.8 mm^3^ for cortical and subcortical targets respectively. This compares to measured averages of 57.5 ± 42.0 mm^3^ and 184.0 ± 82.2 mm^3^ for cortical and deep targets, respectively.

**Figure 6.**
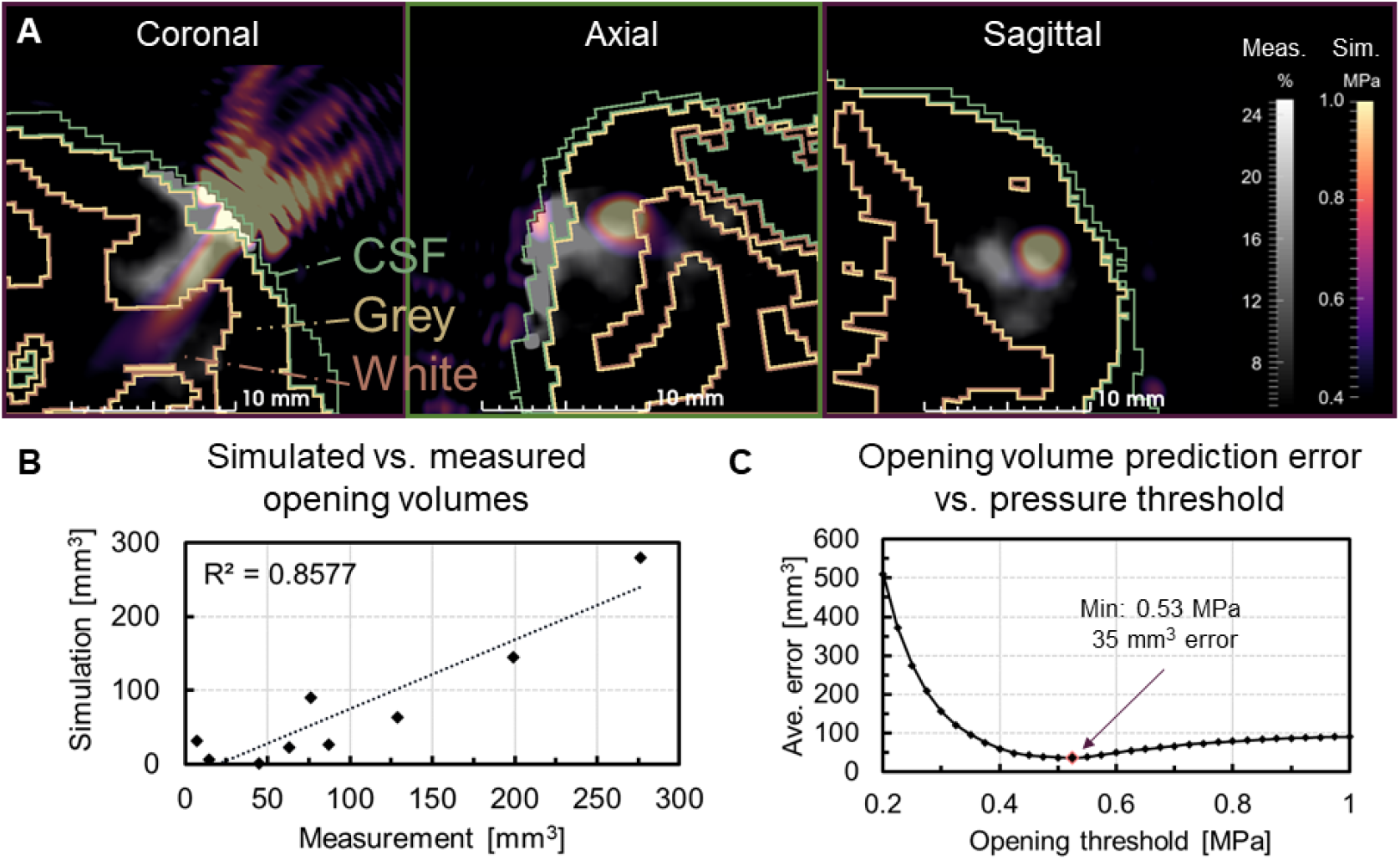
Simulating conditions from in vivo therapies. A) Simulated pressure fields (magma colormap) overlaid on measured percent change images (grayscale colormap) and on tissue type segmentation. Coronal, axial, and sagittal slices are shown. B) Simulated opening volumes plotted against measured opening volumes. C) Opening volume prediction error versus pressure threshold applied to simulated pressure maps. Meas. (Measured), Sim. (Simulated), Max (Maximum), Min (Minimum), CSF (Cerebral Spinal Fluid).

### Safety evaluation

Of the twelve therapies, four exhibited temporary FLAIR hyperintensity at or near the same region as BBBO with no SWI darkening (figure 7). Temporary FLAIR hyperintensity measured at short time delays following a therapy suggests edema occurred in these cases (Ho, Rojas, and Eisenberg 2012). One of the twelve therapies displayed permanent, localized SWI darkening and temporary FLAIR hyperintensity at the target indicating that both edema and extravasation of red blood cells (RBC) occurred. This case occurred in a subcortical target (caudate) at the highest pressure tested (1.4 MPa mean PNP, 1.6 MPa max PNP). Figure 5 distinguishes the datapoints corresponding to edema and/or RBC extravasation with yellow and red markers. The lowest pressure therapy which resulted in temporary edema was at a mean PNP of 0.7 MPa and a maximum PNP of 0.9 MPa. The other four cases which showed edema account for the four highest mean and maximum PNPs tested (1.0 to 1.4 MPa mean, 1.3 to 1.6 MPa maximum). The SWI and FLAIR images from the high-pressure caudate therapy with both edema and RBC extravasation are displayed in figure 7 with the target region highlighted by red crosshairs. All FLAIR hyperintensities were temporary with no hyperintense region persisting and displaying in the following scan. The corresponding minimum time between adjacent therapies in the same monkey was 3 weeks.

**Figure 7.**
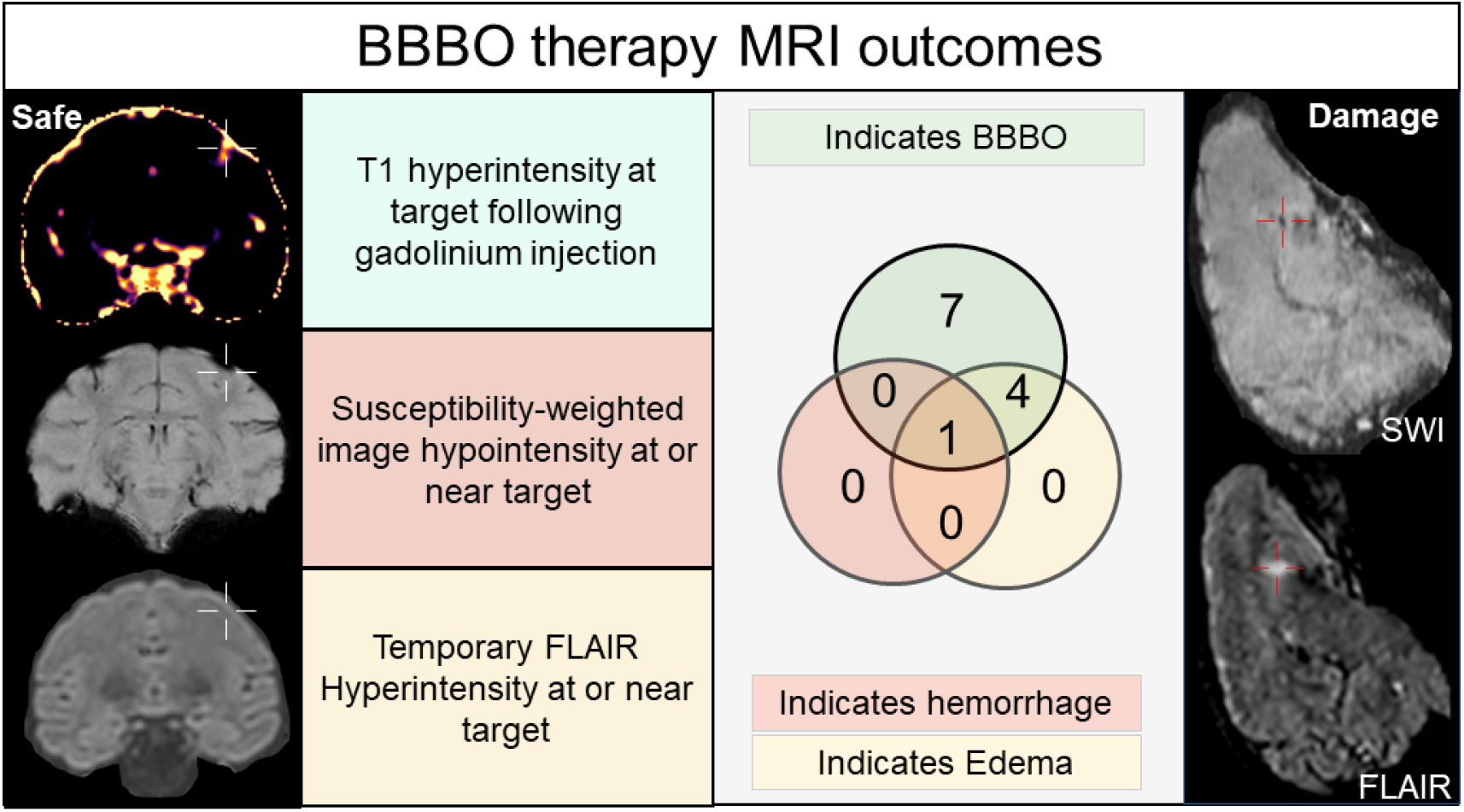
A breakdown of the MR contrasts used during therapies. T1-weighted images following gadolinium injection are sensitive to BBB opening. SWI images are sensitive to hemorrhage. FLAIR is sensitive to edema. Images on the left show one of the seven cases of opening where no SWI or FLAIR abnormalities were visible. The images on the right show the SWI and FLAIR images from the therapy which resulted in both edema and hemorrhage.

### Cavitation monitoring tank measurements

The cavitation monitoring system was developed and tested in a water tank environment using a flow phantom setup with circulating microbubbles matching estimated *in vivo* bubble concentration in the blood. With no skull present, instantaneous stable cavitation dose (SCD) follows a roughly sigmoidal shape showing first increase around 0.35 MPa, rapidly increasing up to 0.7 MPa, and then leveling off at higher pressures (Figure 8). Inertial cavitation dose (ICD) increases starting at 0.5 MPa. Figure 8 (panel A) shows instantaneous cavitation doses with and without a degassed monkey skull present with pressures adjusted for skull transmission (35% transmission for this target). With the skull present, the cavitation signals are largely attenuated. SCD levels at 0.6 MPa decrease from 22,090 in water only to 1,558 with the skull (93% decrease). An increase in SCD is observed from 0.5 to 0.8 MPa. ICD does not increase with increased pressure, despite being in pressure ranges known to produce inertial cavitation.

**Figure 8.**
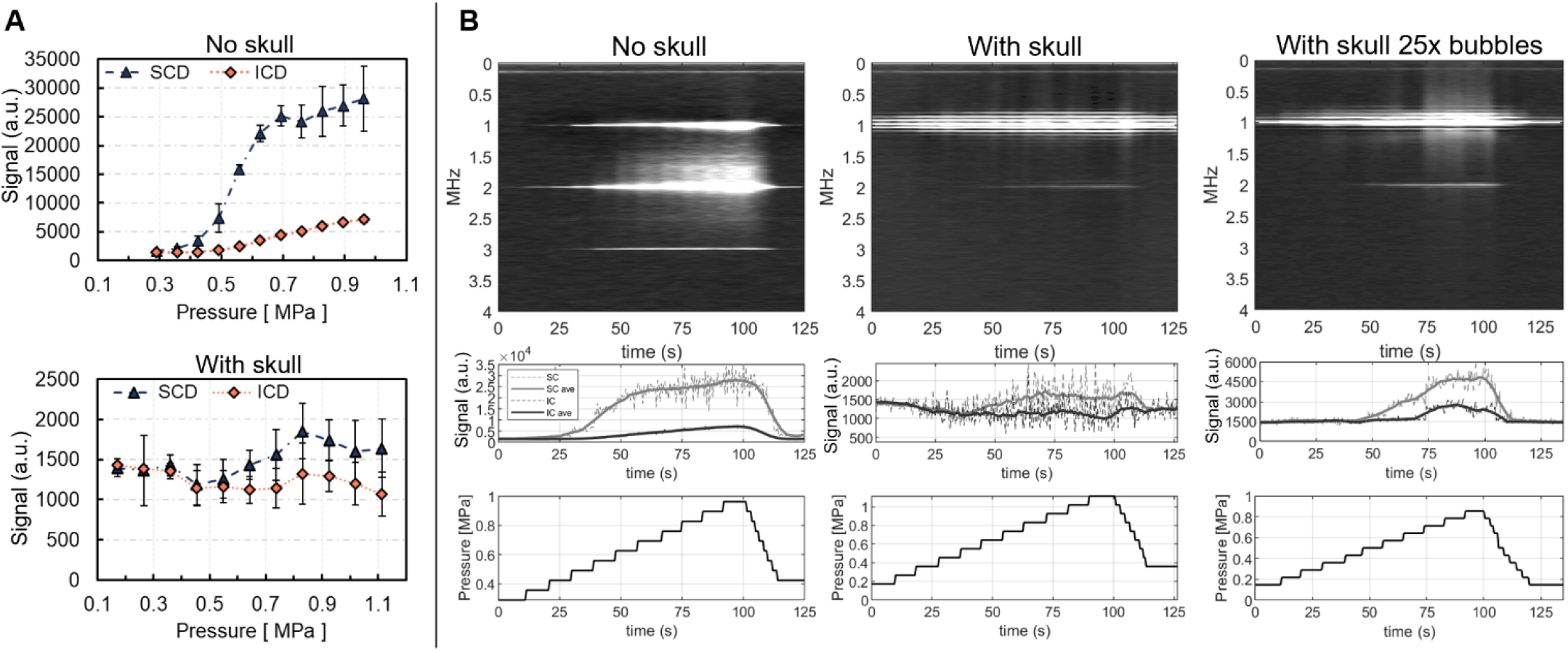
Cavitation measurements made in a microbubble flow phantom. A) Average cavitation signal plotted versus pressure for measurements made with and without an NHP skull in the beam path. B) Cavitation monitoring readouts with for no skull, with skull, and with skull + 25 times higher microbubble concentration than in vivo. The top row shows spectrograms; the middle row show plots of stable (SC) and inertial (IC) cavitation metrics vs. time; the bottom row shows pressure vs. time. Pressures through the skull fragment were adjusted based on 35% transmission.

Figure 8 (panel B) shows cavitation monitoring readouts for water tank measurements with no skull, with a skull, and with a skull using 25 times the estimated *in vivo* concentration of microbubbles. The top row of panel B are spectrograms showing dynamic baseline subtracted spectral content versus time. The second row shows stable cavitation and inertial cavitation metrics plotted with time. The dashed lines are the raw values, the bold lines are averaged with a ten-sample sliding window. The bottom row of panel B shows pressure versus time. Without the skull, stable cavitation is visible in the spectrogram and apparent in the cavitation signal plot beginning at 0.35 MPa and increases with pressure. Inertial signal becomes visible in the spectrogram and cavitation dose plot beginning at 0.50 MPa and increases with pressure. With the skull, inertial cavitation signal is not visualized in the spectrogram nor apparent in the cavitation signal plots. Artifacts are visible around the fundamental frequency band. Some stable cavitation signal is apparent starting at 0.55 MPa.

Increasing the bubble concentration to 25 times the *in vivo* concentration has several effects on the cavitation readouts through the skull. The SCD amplitude increases by a factor of 3 approximately. ICD becomes clearly visible around 1.2 MHz in the spectrogram and in the cavitation signal plots starting at 0.70 MPa. The broadband ICD signal is concentrated at lower frequencies than in the no skull case likely due to frequency dependent attenuation.

### Cavitation monitoring in vivo measurements

All BBBO therapies incorporated real-time cavitation monitoring facilitated by our custom software. Figure 9 shows a representation of the full range of data with four cases showing clear cavitation signal (panel A) and four cases with low cavitation signal readout (panel B) presented to display the range in signal qualities. The opening volume along with any adverse effects are shown above the accompanying data group. The main distinguishing factor between the panel A and panel B is that the panel A groups show clear signal changes in the spectrogram and cavitation metric plots which temporally follow changes in applied pressure shown at the bottom of each group. In panel B, the spectrogram and cavitation plots are unresponsive to changes in pressure. In all therapies, bubbles are slowly injected at the start of the therapy and arrive between 15 and 45 seconds. This arrival is visible in several of the spectrograms and cavitation metric plots.

**Figure 9.**
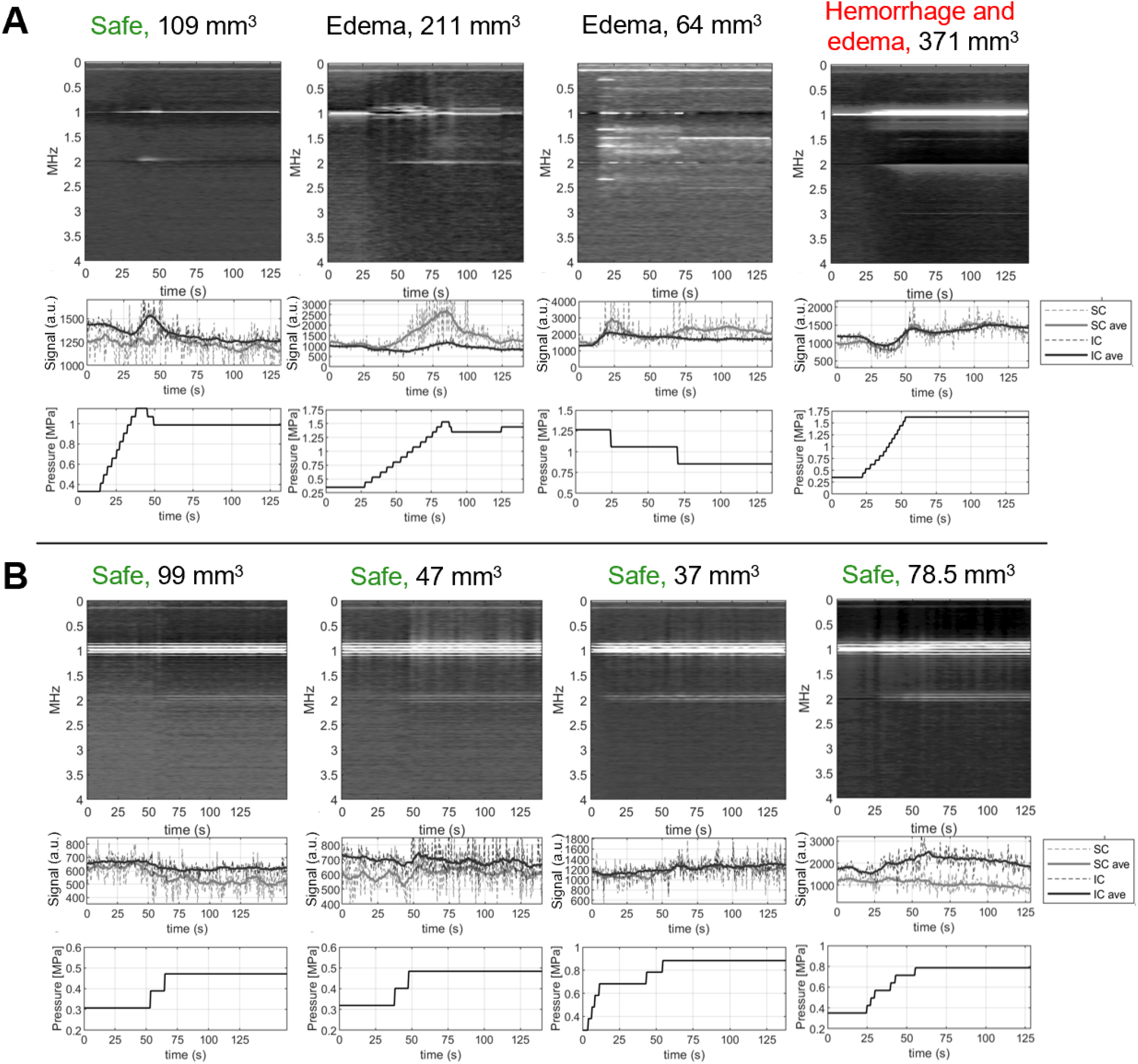
In vivo cavitation monitoring data from eight therapies. Above each therapy are outcomes based on safety scans and opening volume. A) Four therapies where cavitation signals change with pressure and help inform in situ pressure levels. B) Four lower pressure therapies where cavitation signals did not change with pressure and do not inform in situ pressure levels.

In figure 9 panel A first therapy, we ramped the pressure from 0.3 to 1.1 MPa. SCD did not increase before ICD in this case. Seeing ICD increase, we reduced the pressure to 1 MPa. This therapy resulted in neither edema nor RBC extravasation. In Panel A 2^nd^ therapy, we also ramped the pressure. As we ramped, we first noted an increase in SCD between 0.3 to 1.5 MPa. At the upper end of the pressure ramp, we noted ICD and reduced the pressure. This therapy resulted in temporary edema in a small cortical region above the target (putamen, subcortical). In panel A 3^rd^ therapy, clear ultra-harmonics can be noted at the arrival of bubbles which occurred around 15 s into the therapy. Noting the presence increase in ICD also, we dropped the pressure until only SCD was visible. This case resulted in temporary edema. This case occurred early in experiment order and highlighted the need to ramp pressure starting at lower pressure (∼0.4 MPa). For the rightmost case which generated edema and hemorrhage, we ramped the pressure from 0.4 to 1.6 MPa. ICD and SCD both increase starting at 1 MPa. The amplitude of this change was small and was not visible at the time of the therapy due to ineffective window and leveling in the spectrogram. As a result, the therapy pressure was left at a high value for the remainder of the procedure.

Panel B displays results from four lower pressure therapies with maximum pressures ranging between 0.5 MPa and 0.9 MPa. In these cases, the spectrograms and cavitation dose plots are largely unresponsive with changes in pressure. Artifacts seen in the spectrograms as horizontal lines are a product of the subtraction of baseline spectrograms combined with tight colormap windows approaching the noise floor. Despite having no discernable cavitation readout, all four of these therapies resulted in measurable gadolinium perfusion into regions around the target, indicating opening with no adverse effects indicated by SWI or FLAIR images. Figure 10 shows stable and inertial cavitation dose compared with mean therapy pressure and opening volume. Stable cavitation is correlated with mean therapy pressure (R^2^ = 0.454). Other metrics are uncorrelated.

**Figure 10.**
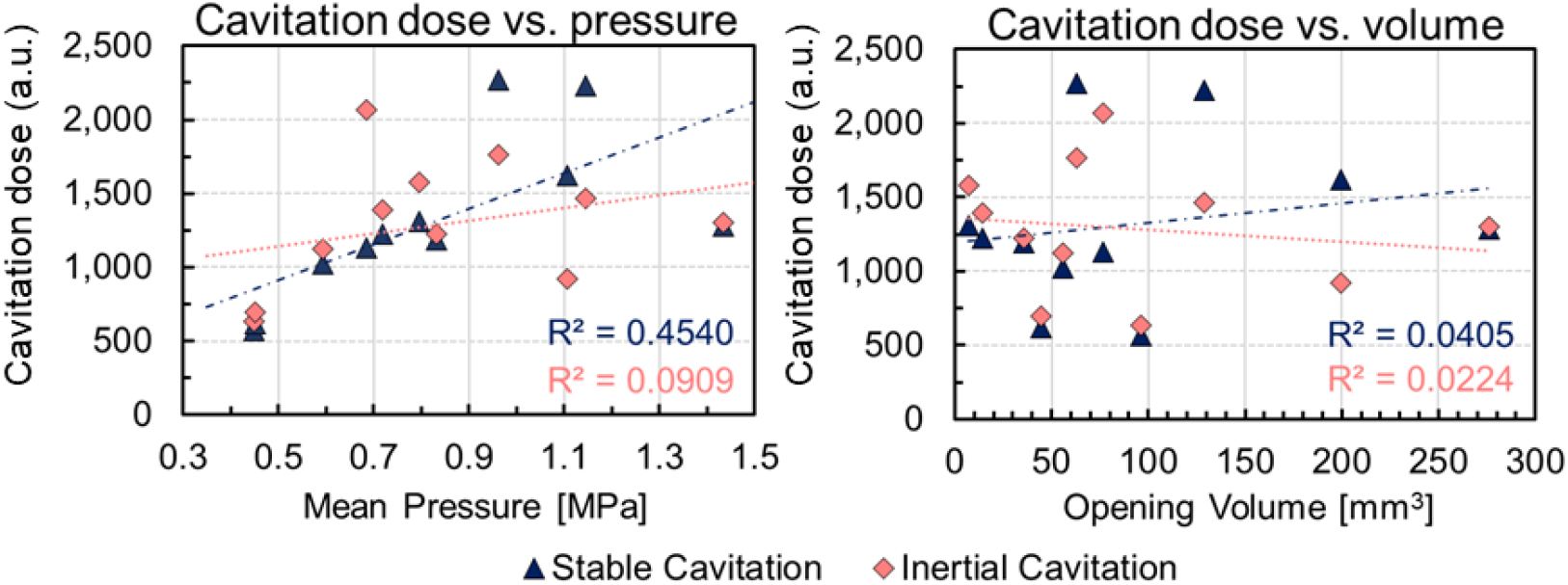
Cavitation dose compared to pressure and opening volume. Stable cavitation dose was partially correlated (R2 = 0.454) with mean therapy pressure. All other comparisons were uncorrelated.

## Discussion

FUS mediated BBB opening has emerged as a critical tool for delivery of therapeutics to the brain. Thus far studies have measured the safety of FUS BBB opening and the size and range of therapeutics able to transport across the permeabilized vasculature (Cammalleri et al. 2020; Song, Harvey, and Borden 2018). The volume of FUS BBB opening determines the extent of the therapy’s effect. It is therefore desirable to achieve small focal spot sizes in applications that target small brain regions. In this work we tested a transducer optimized for small volume BBB opening in the macaque (Manuel, Phipps, and Caskey 2022) intended for gene therapy at the FEF to enable acoustically targeted chemogenetics (Szablowski et al. 2018). The transducer was successful in achieving small volume BBB opening with an average opening volume of 59 ± 37 mm^3^ and 184 ± 2 mm^3^ in cortical and subcortical targets respectively. Other studies in macaques report much larger opening volumes: (NHP 1 462.0 ± 93.4 mm^3^, NHP 2 605.3 ± 253.2 mm^3^ (Downs, Buch, Karakatsani, et al. 2015); 142 mm^3^ to 854 mm^3^ (Karakatsani et al. 2017); 100 to 600 mm^3^ (Wu et al. 2018); 680 mm^3^ to 1413 mm^3^ (Pouliopoulos et al. 2021). By demonstrating that smaller volume opening is possible in macaques we have improved the spatial selectivity of FUS BBB opening to smaller targets. Our system produced larger openings than a recent system tested in a porcine model which produced opening volumes from 3.8 to 53.6 mm^3^ with a 500 kHz, 0.8 f-number transducer (Chien, Xu, et al. 2022).

While the variance of our measured opening volumes is large, simulations accounted for a large portion of this variance (R^2^ = 0.8577). This suggests that the 85% of the variance can be attributed to factors which are captured by the simulation. These factors include *in situ* pressure which is influenced by the transmitted pressure amplitude, the angle of the skull relative to beam propagation, and local skull properties. These observations are consistent with prior works which have demonstrated a dependence of opening volume on pressure (Chen and Konofagou 2014) and targeting effects (Karakatsani et al. 2017) and align with prior work which compared simulations with measured BBB opening outcomes (Wu et al. 2018). Simulations also incorporate the overlap of grey matter with the acoustic focus by including tissue segmentation in the analysis pipeline to account for the higher sensitivity of grey matter to FUS BBB opening. Running these simulations preoperatively could provide a means to minimize variance from trial to trial by adjusting applied pressure based on simulated opening volumes. MR-ARFI may also provide a means for accounting for variance from acoustic scenarios by sampling displacement prior to each therapy (N. Li et al. 2022).

### Grey matter vs white matter opening

Most measured opening occurred in grey matter with 92% of opening in grey at cortical targets and 84% at subcortical targets. Smaller increases in signal were measurable in white matter contiguous with grey matter opening sites. McDannold et al. found opening in white matter of macaques was not visible with gadolinium imaging but was apparent with Evans blue dye (N. McDannold et al. 2012). On the other hand, (Karakatsani et al. 2017) found 80% of measured opening to be in grey matter and 20% in white matter. Across our results and these two studies, grey matter signal increase is brighter than white matter signal increase. This is attributed to the vascular differences in grey and white matter with grey matter being more perfused. Opening in white matter has been reported in humans in an Alzheimer’s related clinical trial (Lipsman et al. 2018) and exclusively at the highest pressures studied in a glioblastoma clinical trial (Carpentier et al. 2016).

### Concentration and delivery efficiency estimation

The concentration of gadolinium at target regions and the delivery efficiency of FUS BBBO for gadolinium can be estimated from the magnitude of signal change after gadolinium administration (Marty et al. 2012). A similar analysis has been conducted by Samiotaki et. al with multiple 5 different flip angle T1-weighted images (Samiotaki et al. 2017). Here we report the same metrics but using a single pre and post therapy T1-weighted image. From the Bloch equations, the longitudinal relaxation signal (*S*) is (Bloch 1946):

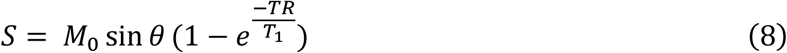

Where *M*_0_ is the equilibrium magnetization, θis the pulse flip angle in degrees, *TR* is the repetition time, and *T*_1_ is the tissue specific longitudinal relaxation time (1.33 s for macaque grey matter at 3 tesla)(Wright et al. 2008). For our *T*_1_ weighted images, *TR*≪*T*_1_, so 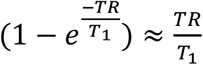. Factoring this in and using relaxation rate 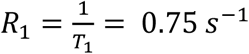, (8) simplifies to:

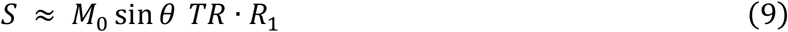

Before gadolinium (injected in the form of Gadavist), the relaxation rate is *R*_1_. After gadolinium, it becomes *R*_1_ + *k[Gd*_*brain*_], where *k* is the relaxivity of gadolinium at 3 Tesla (5.0 L mmol^-1^s^-1^) (Rohrer et al. 2005) and *[Gd*_*brain*_] is the concentration of gadolinium in the image voxel in mmol L^-1^. The fractional change in signal from the pre gadolinium scan *S*_1_ to the post gadolinium scan *S*_2_is given by:

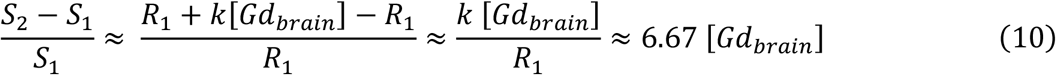

From (10), 10%, 20%, and 30% change in signal in an image voxel correspond to *[Gd*_*brain*_] concentrations of .015 mmol L^-1^, .030 mmol L^-1^, and .045 mmol L^-1^. For any given therapy, this concentration can be converted to amount of gadolinium by voxel-wise multiplication of the voxel volume in L. Using 10% change as a threshold, and calculating delivery on a voxel-wise basis (% change signals range from 10%-80% within voxels), our largest opening (371 mm^3^) delivered a total of 1.05E-5 mmol of gadolinium. Gadolinium was administered at 0.1 mmol kg^-1^, or 1.1 mmol total for an 11 kg monkey. For this therapy, the fraction of administered gadolinium delivered to the target was 0.00095 %. The smallest opening for this macaque (8 mm^3^) delivered 1.52E-7 mmol of Gadavist which corresponds to .000014 % of the administered dose.

It is also possible to estimate the volume of plasma leaked into the extravascular space of the brain. The blood volume (BV) of rhesus macaques is around 60 mL kg^-1^ (Bender 1955), and the extracellular fluid volume (ECF) is approximately 250 ± 40.7 mL kg^-1^ (Overman and Feldman 1947). The injected gadolinium distributes into most tissues other than brain which can be approximated as the sum of the BV and ECF, or 310 ml kg^-1^. The concentration of gadolinium in the blood at the time of injection is 0.1 mmol kg^-1^ diluted into 60 ml kg^-1^ blood, but at the time of BBB opening, this concentration has diluted to 0.1 mmol per 310 ml (.00032 mmol ml^-1^). Because (10) allows us to measure the mmol of gadolinium delivered into the brain, we can estimate the volume of plasma leaked across the BBB as:

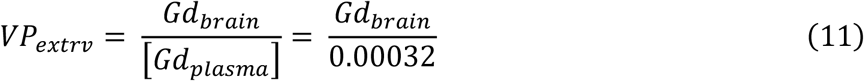

Here *VP*_*extrv*_is the volume of blood extravasated in mL, *Gd*_*brain*_is the amount of gadolinium leaked into the brain in mmol, *[Gd*_*plasma*_] is the concentration of gadolinium in the plasma at the time of BBB opening (0.00032 mmol mL^-1^ for all therapies). For the largest opening, 1.05E-5 mmol of gadolinium was delivered which corresponds to 0.033 mL of plasma. For the smallest opening, 1.52E-7 mmol of gadolinium was delivered or 0.00048 mL of plasma.

### Challenges due to high frequency

The most similar systematic study of BBB opening parameters at 1 MHz (excluding small animals) is the work by Carpentier et. al, who quantified opening outcomes with a 1 MHz implant in humans at pressures from 0.5 to 1.1 MPa (Carpentier et al. 2016). 1 out of 11 patients demonstrated opening in grey matter at 0.8 MPa compared to 3 out of 7 at 0.95 MPa and 12 out of 14 at 1.1 MPa. We tested a pressure range which spanned these values plus additional range. Once hemorrhage was detected at an estimated mean pressure of 1.4 MPa with our system, no pressures at or above that range were tested again.

While the 1 MHz frequency allowed for a small transmit focus to be created, reception of echoes through the skull was challenging due to frequency-dependent attenuation. We identified multiple pressures where ultrasound induced BBB opening but did not generate significant increases in stable cavitation from baseline, making feedback difficult. Work by Wu et. al. investigated the effects of macaque and human skull on a cavitation monitoring system with a 500 kHz transmit frequency (Wu et al. 2014). They noted that the presence of skull increased the pressure detection threshold for ultra-harmonic and inertial cavitation signals. We noted similar trends with our system in flow phantom measurements, and the effect will likely be more present in our system due to our higher transmit frequency. Additionally, the skull produces strong reflections which impinge on the PCD during cavitation measurements. Increasing the transmit amplitude increases the amplitude of these reflections which have inherent harmonic content even in the absence of bubbles. Following methodologies proposed by Kamimura et. al we employed baseline subtractions using echoes acquired prior to microbubble administration (H. A. Kamimura et al. 2019). In the absence of baseline subtraction, ICD and SCD increase with increasing pressure even in the absence of microbubbles, presumably due to non-linear echoes from the skull. Although baseline subtraction helped isolate echoes arising from microbubbles, we were not able to detect all bubble activity.

### Cavitation monitoring limitations

While our cavitation monitoring provided useful readouts for several therapies, when combined across trials cavitation doses did not correlate with opening volume or safety outcomes for our system. In a similar study with a 500 kHz transducer in macaques, cavitation metrics also did not correlate with opening volume (Marquet et al. 2014). In several cases shown in figure 9, there are clear signatures of harmonic and broadband signals. However, the amplitude and characteristics of these features vary greatly from one therapy to another and are mostly absent at therapies which resulted in neither edema nor hemorrhage. A possible approach for therapy monitoring in future uses may be to ramp pressure up to 0.7 MPa *in situ*, at which point the therapy can progress so long as no harmonic or inertial signals are present. If either harmonic or inertial signals are present, this likely suggests higher than normal transmission and pressure should be reduced until the signals are absent.

The geometric arrangement and frequency response of the cavitation monitor are known to be factors in sensitivity and could be improved upon to yield greater monitoring. A recent study in a porcine model achieved effective cavitation monitoring by tracking cavitation at the fourth harmonic of a 500 kHz transducer (2 MHz) (Chien, Xu, et al. 2022). This approach used “dummy” signal captures following microbubble injection transmitted at 0.3 MPa to establish a baseline signal level at the fourth harmonic and tuned therapy pressures to increase the same signal metric relative to the baseline (Chien, Yang, et al. 2022). The results from this group indicate that closed-loop feedback is possible in large animal models at the frequency ranges monitored by our system, and suggest that the receive element used on our array lacks sufficient sensitivity. Given the extent of skull attenuation on our cavitation signals, it is likely that incorporating a receive element at the subharmonic (500 kHz) where attenuation is lower could yield better consistency with *in vivo* cavitation monitoring readouts in the presence of thick skulls. Subharmonics have been used successfully in mice for BBB opening cavitation monitoring (Burgess et al. 2014).

### Safety measurements

We detected temporary edema in 5 of 12 cases and hemorrhage in 1 of 12 cases. Temporary edema has been observed by others at lower pressures than hemorrhage and typically resolves within a week. Four instances of edema were measured in a safety study using 500 kHz between 200 and 400 kPa (Downs, Buch, Sierra, et al. 2015). Additionally, four instances of edema were reported at 500 kHz 300 kPa (Downs, Buch, Karakatsani, et al. 2015). Both hemorrhage and edema were observed at 325 kPa, 500 kHz (H. A. Kamimura et al. 2019). In our case, four of the five cases of edema were observed at the four highest mean pressures applied (ranging from 1.0 to 1.4 MPa) which suggest using only lower pressures may avoid edema. However, there was a case of edema at a mean pressure of 0.7 MPa, max pressure 0.9 MPa. Rather than declare a hard threshold for edema, we note that at mean pressures below 0.9 MPa the instances of edema were 1/7 and fell to 0/4 below 0.7 MPa.

### Limitations of the study

Because our therapies were designed to dynamically set pressure based on cavitation signals, *in situ* pressure was not controlled for. It may be the case that opening volumes would have lower variance if pressure were held constant across therapies. However, applying a constant transmitted pressure would not result in constant *in situ* pressure as this will vary from target to target and subject to subject due to differences in the acoustic environment. In addition to varying skull geometries and acoustic properties, internal reflections likely give rise to complex pressure fields through the brain via standing wave formation given our long pulse length (10 ms) (Tang and Clement 2010; O’Reilly, Huang, and Hynynen 2010). Standing waves result in nodes of zero pressure and twice the pressure and may cause opening and/or damage despite low transmit amplitude. The distribution of microbubbles supplied by the vasculature is inhomogeneous in the brain which may partially decouple opening, edema, and hemorrhage from the spatial pressure field (Prada et al. 2021). For this reason, small changes in focus location alone may result in changes in opening volume and cavitation readout. Cavitation monitoring was implemented with the hope of accounting for these challenges but provided insufficient signal in several cases. Future FUS BBB opening transducer designs could prioritize receive sensitivity to address these limitations.

## Conclusion

We tested a transducer optimized for small volume BBB opening in cortical targets in macaques and achieved opening at smaller volumes than previously reported. We characterized opening volume and safety outcomes across a range of pressures and targets, contributing new insight into BBB opening at 1 MHz in macaques. Cavitation monitoring with our 2 MHz receive element provided insight during some high-pressure therapies but had low SNR at lower pressure levels. This system improves FUS BBB opening spatial specificity in macaques.

## Acknowledgments

NIH UG3 MH120102 and S10ODO21771-01 for funding.

